# Rotavirus NSP1 localizes in the nucleus to disrupt PML nuclear bodies during infection

**DOI:** 10.1101/619932

**Authors:** Samantha K. Murphy, Michelle M. Arnold

## Abstract

The rotavirus nonstructural protein 1 (NSP1) antagonizes interferon (IFN) induction in infected host cells. The primary function of NSP1 is thought to be degradation of interferon regulatory factors (IRFs) and beta-transducin repeat-containing protein (β-TrCP) in the cytoplasm to inhibit IFN induction. Here, we report that NSP1 localizes to the cytoplasm and nucleus and disrupts promyelocytic (PML) nuclear bodies (NB) in the nucleus during infection. Nuclear localization of NSP1 did not require an intact C terminus, suggesting NSP1 has a novel function in the nucleus independent of degradation of IRFs or β-TrCP. NSP1 expression either led to a reduction in PML NB number or a change in PML NB morphology from sphere-shaped foci to oblong-shaped structures, depending on the virus strain. Additionally, infection was not affected when cells lack PML NB, suggesting that rotavirus does not require PML for replication in highly permissive cell types. PML was not essential for nuclear localization of NSP1, but PML was required for NSP1 nuclear focus formation. PML NBs play an important role in many cellular functions that include IFN induction and host stress responses. This is the first report that rotavirus, a cytoplasmically replicating virus, encodes a viral protein that localizes to the nucleus during infection, and may suggest a new function of NSP1 in the nucleus.

**IMPORTANCE:** Rotavirus causes severe gastroenteritis in young children and leads to over 200,000 deaths per year. Rotavirus is a cytoplasmically replicating virus, and must find ways to avoid or actively inhibit host antiviral responses to efficiently replicate. The nonstructural protein NSP1 is known to inhibit IFN induction by promoting degradation of host proteins in the cytoplasm of infected cells. Here, we demonstrate that NSP1 also localizes to the nucleus of infected cells, specifically to PML NB. NSP1 causes a disruption of PML NB, which may serve as an additional mechanism of IFN inhibition or interfere with other nuclear processes to promote viral replication. A detailed exploration of the manipulation of nuclear processes in cells infected with cytoplasmically replicating viruses will lead to new insights into viral evasion of host responses.

## INTRODUCTION

Rotaviruses are double-stranded RNA viruses containing a segmented genome packaged inside a multi-layered, non-enveloped viral particle (1). Most steps of the rotavirus replication cycle take place in the cytoplasm of infected cells in replication centers known as viroplasms, and final assembly takes place via a process of budding through the endoplasmic reticulum (ER) (2). Although the virus relies on its host to replicate efficiently, it must also find ways to avoid host antiviral responses. As with many viruses, rotaviruses target several different steps of the type I interferon (IFN) response in order to prevent the expression or activity of host antiviral proteins. For instance, the viral protein VP3 functions within the innermost layer of the viral particle as the capping enzyme, but also works within the infected cells to cleave 2’-5’-oligoadenylates to inhibit the activation of ribonuclease L (RNaseL) (3, 4).

Rotavirus also encodes the nonstructural protein NSP1 to inhibit the production of type I IFN, which in turn is thought to limit the expression of antiviral IFN-stimulated genes (ISGs) (5–10). Depending on the virus strain, NSP1 induces proteasomal degradation of IFN regulatory factors (IRFs) or β-TrCP, a protein component of a host cullin-RING E3 ubiquitin ligase complex that activates NF-κB (5, 8, 11, 12). Other cellular targets of NSP1 have also been identified, all of which appear to be targeted for degradation by the proteasome (13–16). NSP1 has been shown to localize diffusely throughout the cytoplasm of infected cells, but appears to be excluded from the viroplasms (17–19). Recent studies using screening approaches to identify host proteins that associate with NSP1 have found evidence for nuclear proteins in NSP1 pull down samples, but much of this data not been validated or investigated in detail (20, 21).

In this study, we demonstrate that rotavirus NSP1 localization is not restricted to the cytoplasm of infected or transfected cells; NSP1 also localizes to the nucleus, which still occurs in the absence of an intact C-terminal domain. Some rotavirus strains, such as SA11-4F, were found diffusely distributed at low levels in the nucleus of infected cells, while other strains, such as OSU, formed distinct nuclear foci. The rotavirus strains that formed nuclear foci were found to colocalize with promyelocytic (PML) nuclear bodies (NB). NSP1 nuclear localization led to changes in the morphology or a reduction in the number of PML NB, depending on the virus strain. In the absence of PML, virus titers were unchanged, but fewer infected cells contained NSP1 nuclear foci, suggesting the irregularly shaped foci are attributed to the presence of PML. Together, the data suggest that NSP1 can localize to the nucleus to disrupt PML NB. This is the first report of a rotavirus protein found to localize to the nucleus, an unexpected finding given that the virus replicates in the cytoplasm of infected cells.

## RESULTS

### NSP1 localizes to the nucleus and the cytoplasm of infected cells

We previously examined host proteins that associate with NSP1 by performing pull down assays followed by mass spectrometry (20). The NSP1 protein from SA11-4F and OSU strains of rotavirus were found to associate with proteins associated with the nuclear pore, including exportin-1 (CRM1), nuclear pore complex protein NUP205, and NUP93. To determine if NSP1 localizes to the nucleus of infected cells, HT29 cells were mock infected or infected with SA11-4F or OSU rotavirus at a MOI of 5 for 8 h, which is when peak expression of NSP1 is expected to occur during infection (22). Cells were harvested and fractionated into whole cell, cytosolic and nuclear fractions, followed by SDS-PAGE. Immunoblotting for SA11-4F NSP1 and OSU NSP1 demonstrated that the NSP1 proteins were localized to both cytosolic and nuclear fractions of infected cells (Fig. 1A). Nuclear fractions did not contain GAPDH, indicating that they were clear from cytosolic contamination. To ensure efficient separation of the endoplasmic reticulum (ER) from the nuclear membrane, calnexin was included as a control. To demonstrate that nuclear localization was not generalizable to other viral proteins, the viroplasm scaffolding protein NSP2 was detected only in the cytoplasmic fraction.

**FIG 1.**
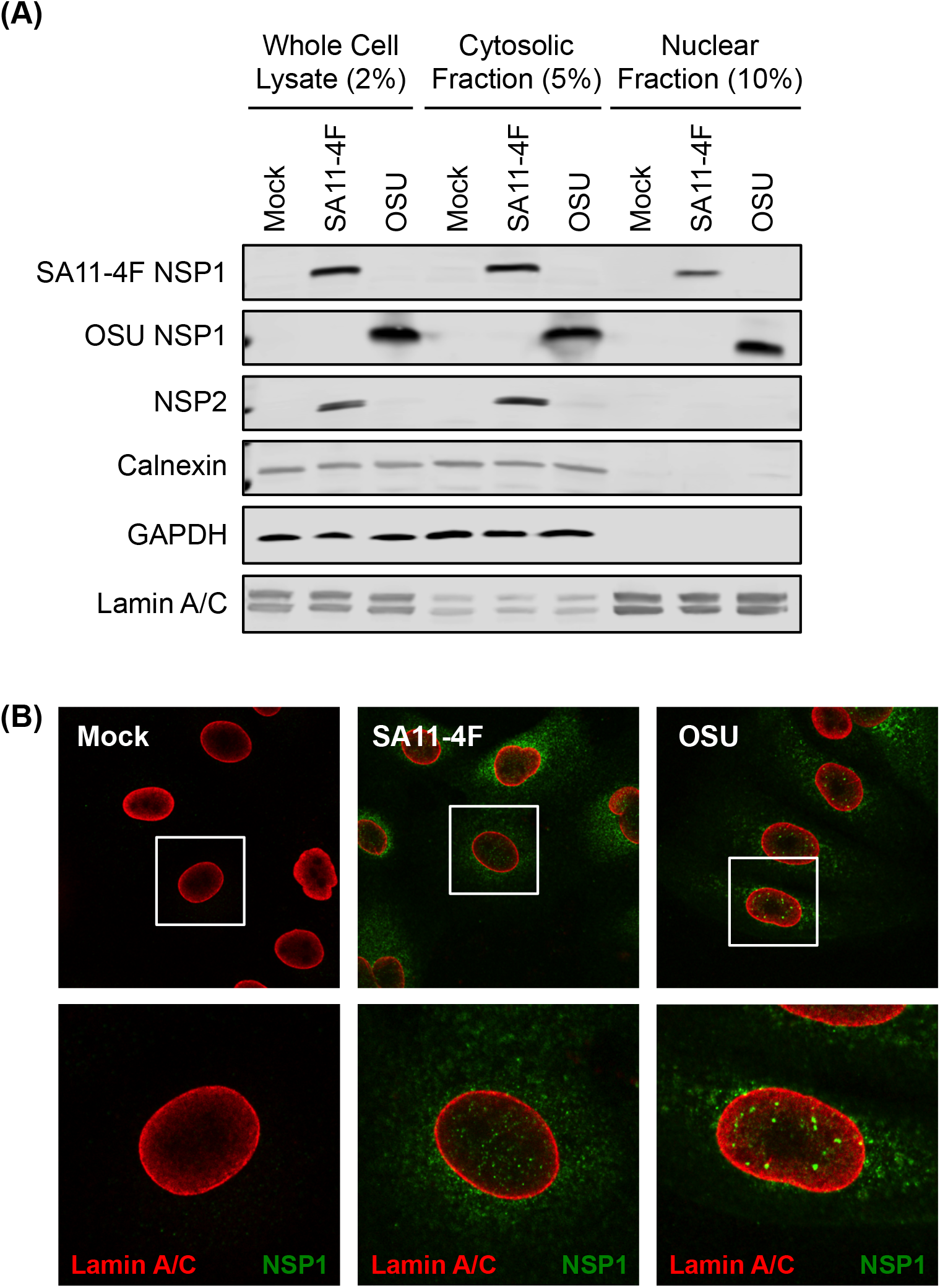
NSP1 localizes to the cytoplasm and nucleus of infected cells. (A) HT29 cells were mock infected or infected with SA11-4F or OSU rotavirus (MOI = 5) for 8 h. Whole cell lysates, cytosolic fractions, and nuclear fractions were resolved by SDS-PAGE and analyzed by immunoblotting for SA11-4F NSP1, OSU NSP1, SA11-4F NSP2, calnexin (loading control to exclude ER contamination), GAPDH (loading control for cytosolic extract), or lamin A/C (loading control for nuclear extract). Blots were imaged using the Odyssey infrared imaging system (LiCor). **(B)** MA104 cells were mock infected or infected with SA11-4F or OSU rotavirus (MOI = 5) for 8 h. Cells were fixed and stained with α-SA11-4F or α-OSU NSP1 (green) and α-lamin A/C (red) antibodies followed by secondary staining with AlexaFluor488 goat α-rabbit and AlexaFluor594 goat α-mouse. Lower panels are zoomed in images of cells in white box in upper panels.

To gain more detailed information about the nuclear distribution of NSP1 in infected cells, MA104 cells were mock infected or infected with SA11-4F or OSU rotavirus at a MOI of 5 for 8 h, followed by immunostaining and confocal microscopy (Fig. 1B). Lamin A/C was used to visualize the boundaries of the nuclear membrane. In cells infected with SA11-4F, NSP1 appeared to be diffuse throughout the cytoplasm and nucleus. However, the nuclear distribution differed in cells infected with OSU, where NSP1 was found to form distinct foci within the nucleus. Together, these data show that NSP1 from SA11-4F and OSU strains of rotavirus is not restricted to the cytoplasm of infected cells, but also localizes to the nucleus.

### OSU-like and UK-like NSP1 proteins form foci within the nucleus

To investigate the nuclear distribution of NSP1 proteins from other strains of rotavirus, the localization of tagged NSP1 proteins from different rotaviruses was examined in transfected cells. A transfection approach was used because antibodies are not available to all NSP1 proteins, and those antibodies that are available tend to not cross-react with NSP1 from different rotavirus isolates (8, 23). 293T cells were transfected with plasmids expressing Halo-tagged NSP1 from SA11-4F, OSU, WI61, UK, and DS-1 rotaviruses and harvested at 48 h p.t. Cells were then fractionated into whole cell, cytosolic and nuclear fractions, followed by SDS-PAGE and immunoblotting. Halo-tagged NSP1 from all strains tested was found to localize to the nuclear fraction of transfected cells (Fig. 2A). The levels of SA11-4F and DS-1 NSP1 proteins in the nucleus were consistently lower than what was observed for OSU, WI61, and UK NSP1 proteins. Although low levels of the HaloTag alone were detectible in the nucleus due to its small size, the levels of Halo-NSP1 proteins were notably higher in the nuclear fraction of transfected cells.

**FIG 2.**
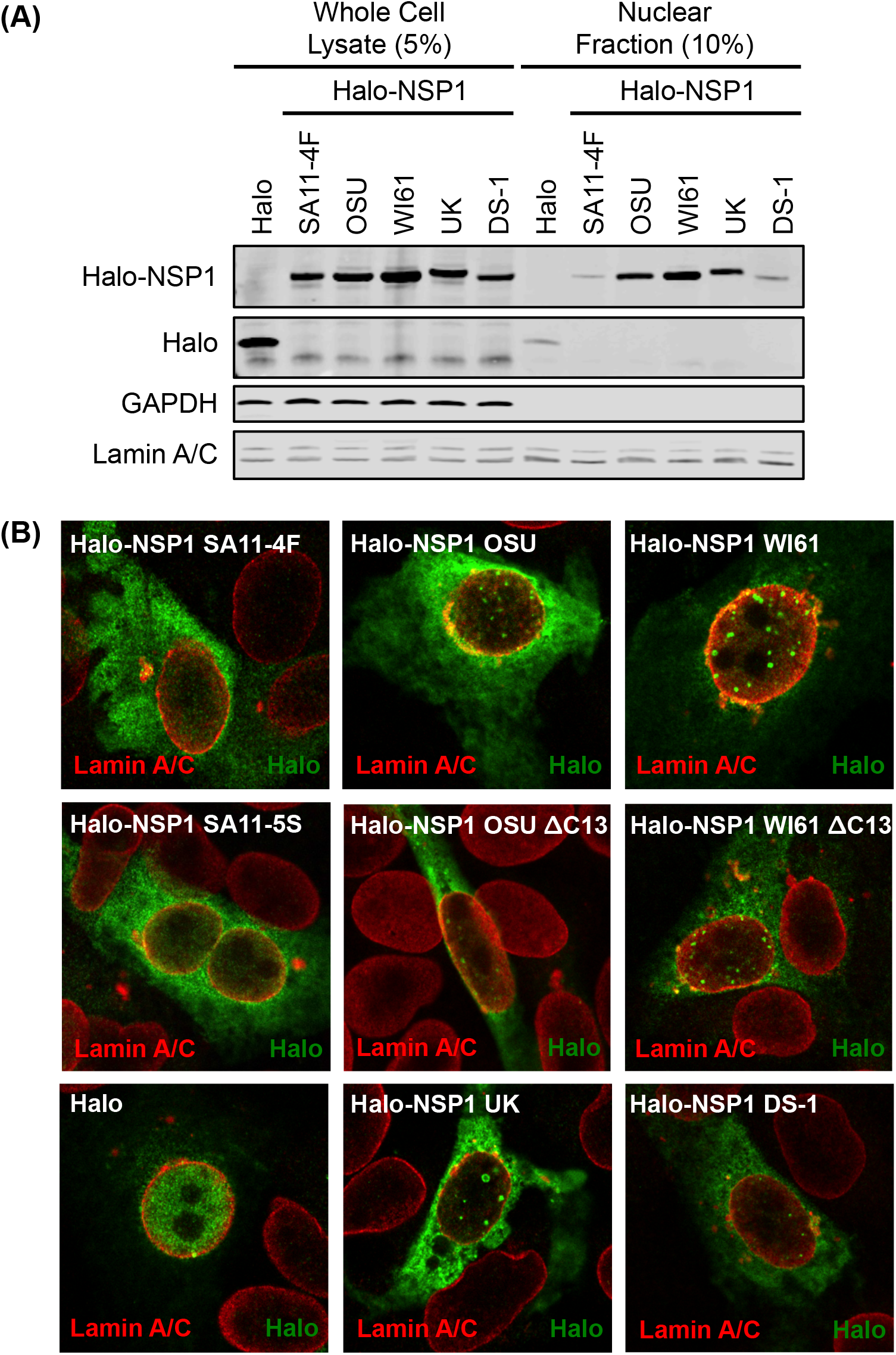
NSP1 localizes to the cytoplasm and nucleus of transfected cells. **(A)** 293T cells were transfected with a plasmid encoding Halo-tagged NSP1 from SA11-4F, OSU, WI61, UK, or DS-1 rotavirus strains, or with empty Halo tag vector, for 48 h. Whole cell lysates, cytosolic fractions, and nuclear fractions were resolved by SDS-PAGE and analyzed by immunoblotting for the Halo tag, GAPDH (loading control for cytosolic fraction), or lamin A/C (loading control for nuclear fraction). Blots were imaged using the Odyssey infrared imaging system (LiCor). **(B)** MA104 cells were transfected with a plasmid encoding Halo-tagged NSP1 from SA11-4F, OSU, WI61, UK, or DS-1 rotavirus strains, small C-terminal NSP1 deletions of SA11-4F (SA11-5S), OSU (OSU ΔC13), WI61 (WI61 ΔC13), or with empty Halo tag vector, for 48 h. Cells were fixed and stained with α-Halo tag (green) and α-lamin A/C (red) antibodies followed by secondary staining with AlexaFluor488 goat α-rabbit and AlexaFluor594 goat α-mouse.

To determine if the subcellular distribution of Halo-tagged NSP1 proteins was similar to that observed in infected cells (Fig. 1B), MA104 cells were transfected with plasmids expressing the same Halo-tagged NSP1s for 48 h, as well as two C-terminally truncated NSP1 proteins, SA11-5S and OSU ΔC13, followed by immunostaining and confocal microscopy (Fig. 2B). Lamin A/C was used to visualize the boundaries of the nuclear membrane. In cells transfected with Halo-NSP1 from SA11-4F rotavirus, the NSP1 protein was diffusely localized throughout the cytoplasm and nucleus, similar to what was observed in infected cells. The SA11-5S NSP1 contains a 17 amino acid C-terminal truncation, but is otherwise identical to SA11-4F NSP1; cells transfected with SA11-5S NSP1 also contained NSP1 in the nucleus in a diffusely distributed pattern. In cells transfected with Halo-NSP1 from OSU, WI61, UK, and DS-1 rotaviruses, the NSP1 protein was diffusely localized throughout the cytoplasm, but distinct NSP1 foci formed in the nucleus. The removal of 13 amino acids from the OSU NSP1 (OSU ΔC13) or WI61 NSP1 (WI61 ΔC13) did not have an impact on the ability of NSP1 to localize and form distinct foci in the nucleus of transfected cells. The levels of nuclear Halo-NSP1 from SA11-4F and DS-1 in immunostained cells appeared to be rather low (Fig. 2B), as was reflected in the fractionation assay (Fig. 2A). Together, these data show that NSP1 from several different strains of rotavirus localizes to both the cytoplasm and nucleus, and that in most cases, excepting the SA11-4F and SA11-5S NSP1, distinct nuclear foci are formed by NSP1.

### OSU NSP1 forms foci in the nucleus at early times post-infection

To determine at what time post-infection NSP1 begins to accumulate in the nucleus of infected cells, MA104 cells were mock infected or infected with SA11-4F or OSU rotavirus and harvested at 2 h intervals for 12 h total. Fixed cells were then immunostained with antibodies specific to SA11-4F NSP1 or OSU NSP1 (8). In mock-infected cells stained with the SA11-4F NSP1 antibody, no signal was detected, suggesting that the SA11-4F NSP1 did not cross-react with any cellular proteins (Fig. 3A). In SA11-4F-infected cells, NSP1 was first observed in the cytoplasm and the nucleus at 6 h p.i., and continued to accumulate through 12 h p.i. SA11-4F NSP1 was diffusely distributed in the nucleus, and did not form distinct foci during the course of infection. At late times post-infection (10 and 12 h p.i.), the cytoplasmic NSP1 appeared to be excluded from the viroplasms, as has previously been observed (17). In mock-infected cells stained with the OSU NSP1 antibody, a low-level background signal was detected, indicating that the OSU NSP1 antibody cross-reacted with a cellular protein (Fig. 3B). However, the accumulation of OSU NSP1 over the background staining was evident by 4 h p.i., and levels of OSU NSP1 continued to increase through 12 h p.i. Notably, the OSU NSP1 nuclear foci appeared by 6 h p.i., suggesting their formation was not due to over-expression of NSP1 protein at late times post-infection.

**FIG 3.**
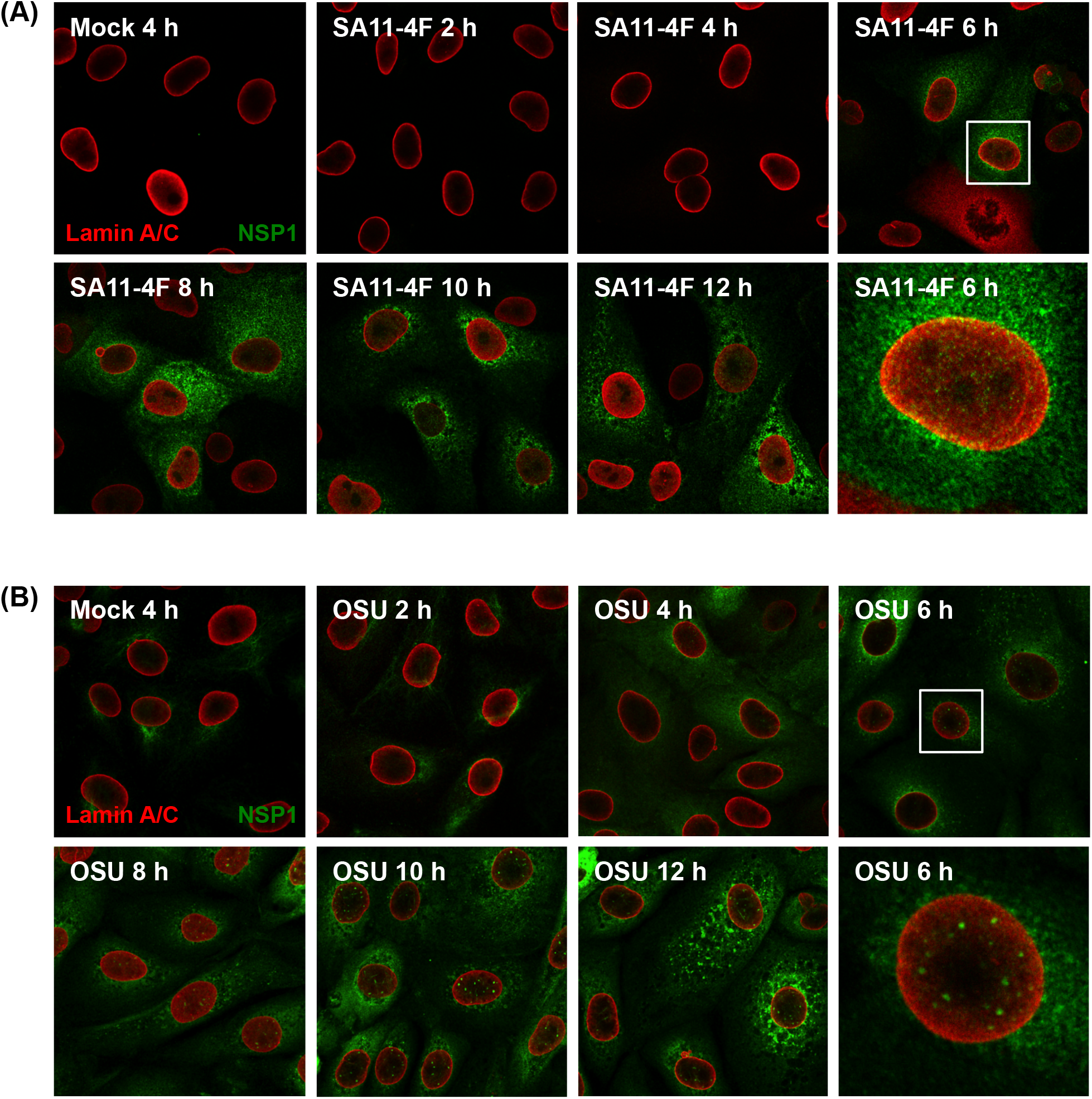
Time course of NSP1 nuclear accumulation in infected cells. MA104 cells were mock infected or infected with SA11-4F or OSU rotavirus (MOI = 5) for 2, 4, 6, 8, 10 or 12 h. Cells were fixed and stained for **(A) α**-SA11-4F NSP1 (green), or **(B)** α-OSU NSP1 (green), and α-lamin A/C (red) antibodies followed by secondary staining with AlexaFluor488 goat α-rabbit and AlexaFluor594 goat α-mouse. A zoomed in image at 6 h p.i. is included for SA11-4F and OSU infected cells, which are surrounded by a white box in upper panels.

### OSU NSP1 nuclear foci colocalize with PML nuclear bodies during infection

To determine the nature of the NSP1 nuclear structures in OSU-infected cells, HaCaT cells were mock infected or infected with SA11-4F or OSU rotavirus and fixed at 8 h p.i.. HaCaT cells are a human keratinocyte cell line that is highly amenable to immunofluorescence microscopy. Cells were co-stained with an antibody to NSP1 and markers of nuclear gems (gemin 2), nuclear speckles (SC35, also known as the serine/arginine-rich splicing factor SRSF2), or promyelocytic leukemia (PML) nuclear bodies (NB). A colocalization histogram was generated for imaged cells to determine if the signals from NSP1 and the subnuclear structures occurred in the same space. In mock-infected cells, gemin 2 appeared as one or two sphere-shaped foci in the nucleus, and no cross-reactivity with the SA11-4F NSP1 antibody was observed (Fig. 4A). In cells infected with SA11-4F or OSU rotavirus, no colocalization was apparent between nuclear NSP1 from either virus and the gemin 2 protein. Additionally, there appeared to be no major changes in the morphology or distribution of nuclear gems in rotavirus infected cells. Nuclear speckles were stained with an antibody to SC35, which appeared as irregularly distributed areas throughout the nucleus of mock-infected cells (Fig. 4B). In cells infected with SA11-4F or OSU rotavirus, no colocalization was apparent between nuclear NSP1 from either virus and SC35, nor was there a change in the morphology and distribution of nuclear speckles in infected cells.

**FIG 4.**
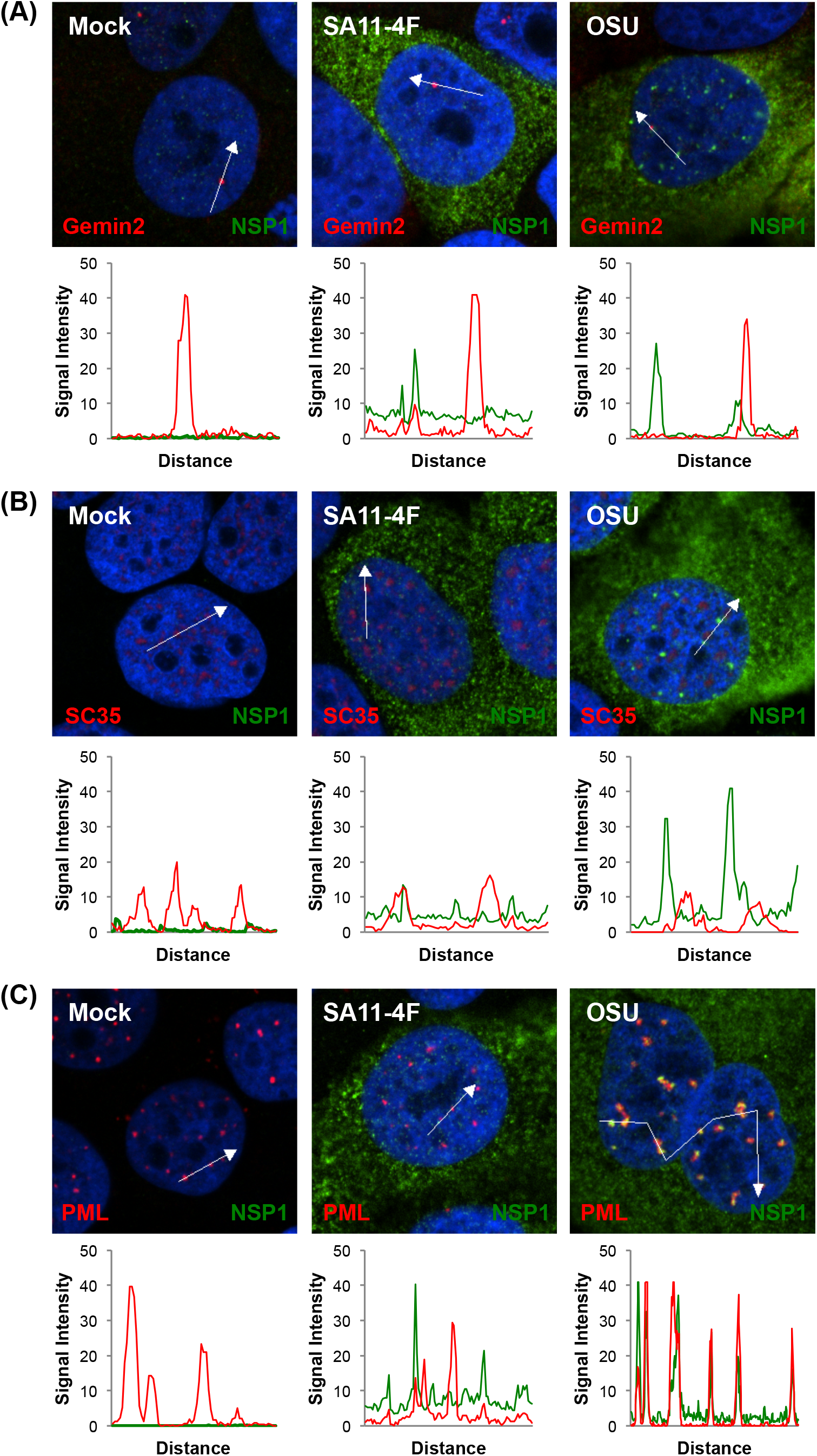
Intranuclear OSU NSP1 localizes to PML nuclear bodies. HaCaT cells were mock-, SA11-4F-, or OSU-infected (MOI = 5) for 8 h. Cells were fixed and stained with α-SA11-4F or α-OSU NSP1 (green) antibodies and **(A)** α-Gemin2 (red), **(B)** α-SC35 (red), or **(C)** α-PML (red) antibodies. Nuclei were counterstained with DAPI (blue). Histograms display measured fluorescence signal intensity (x100) along the arrow in the image panels.

Interestingly, in OSU-infected cells the NSP1 foci localized in the nucleus were found to colocalize with PML NB (Fig. 4C). Visual inspection of immunofluorescence images showed overlap between the signal from the OSU NSP1 and the PML antibody, and the OSU NSP1 and PML signal peaks corresponded with one another in the histogram profile. There also appeared to be changes in the morphology of PML NB during infection with OSU when compared to PML NB in mock-infected cells, where PML staining appeared as sphere-shaped foci throughout the nucleus. Colocalization was not observed between SA11-4F NSP1 and PML in infected cells, likely because of the diffuse distribution pattern of SA11-4F NSP1 in the nucleus. Similar results were observed in infected MA104 cells and MA104 cells transfected with plasmids encoding Halo-tagged NSP1 (data not shown). Together, the data indicate that OSU NSP1 nuclear foci colocalize with PML NB in infected cells.

### Changes in the number and area of PML NB in rotavirus infected cells

To follow up on the observation that the morphology of PML NB appeared to be altered in OSU-infected HaCaT cells, measurements of the number and area of PML NB present in the nucleus of rotavirus infected cells were made in comparison to uninfected cells. MA104 cells were mock infected or infected with SA11-4F, SA11-5S, or OSU rotaviruses and fixed at 8 h p.i. The SA11-5S rotavirus is identical to the SA11-4F virus strain except for a rearrangement in the NSP1 coding gene that results in a small C-terminal deletion in the NSP1 protein; this truncated NSP1 protein is no longer able to bind to IRF3 and target it for degradation (5, 11). Because of this C-terminal deletion, the SA11-5S NSP1 protein is no longer detected by available NSP1 antibodies, thus the cells were stained with an antibody to the viral protein VP6 to detect infected cells, and an antibody to PML to detect the PML NB (Fig. 5A). The number of PML NB in 54 cells was counted in maximum projection of z-stack images of each condition, and the area of 100 PML NB was measured for quantification. In mock-infected cells, PML NB appeared sphere-shaped in the nucleus (Fig. 5A). Mock-infected cells on average contained approximately 12 PML NB per nucleus with an area of about 0.21 μm^2^ (Fig. 5B, 5C).

**FIG 5.**
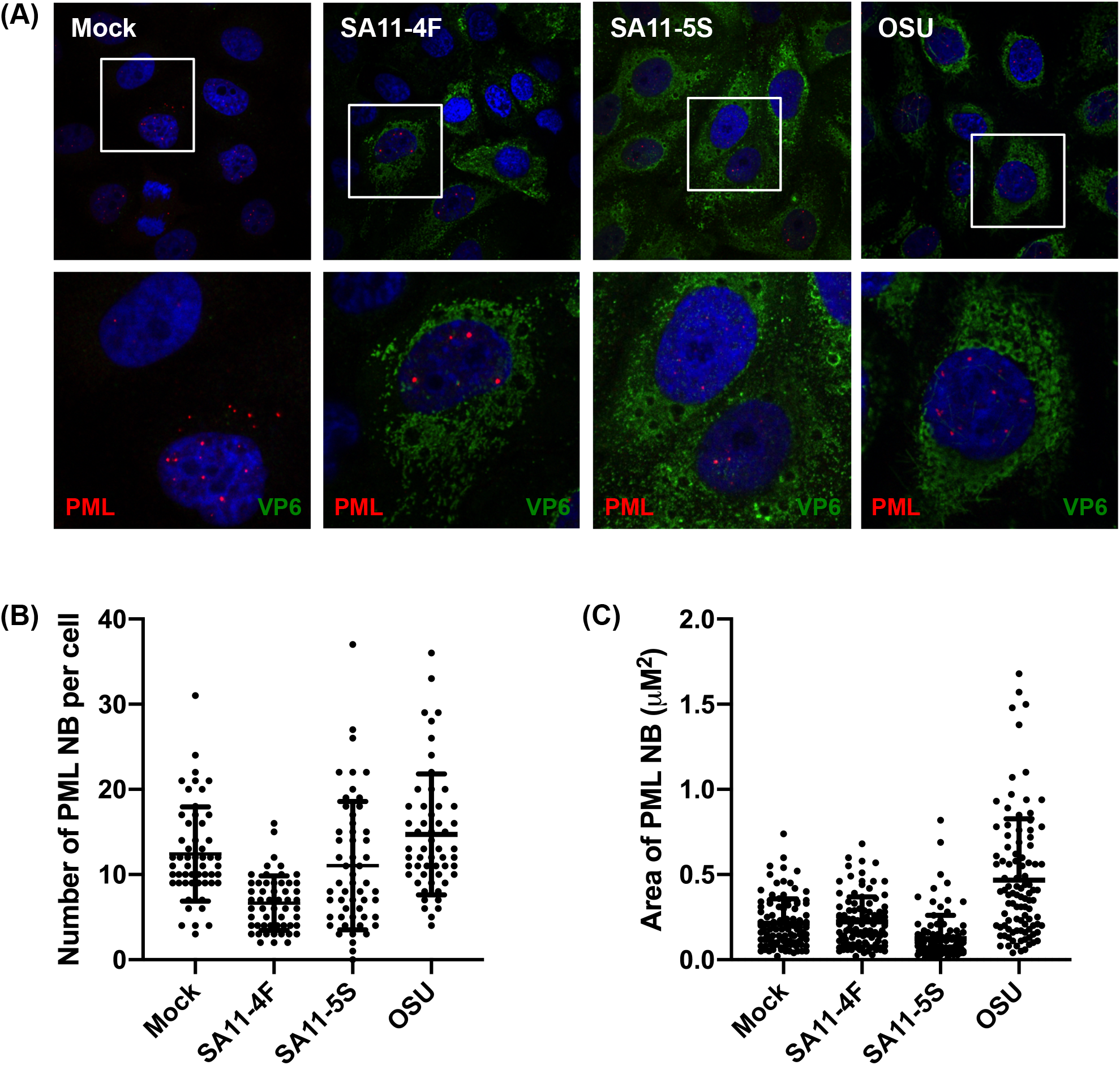
PML body morphology, quantity, and size in rotavirus-infected cells. **(A)** MA104 cells were mock infected or infected with SA11-4F, SA11-5S, or OSU rotavirus (MOI = 5) for 8 h. Cells were fixed and stained with α-VP6 (green) and α-PML (red) antibodies followed by secondary staining with AlexaFluor488 goat α-guinea pig and AlexaFluor594 goat α-mouse. Nuclei were counterstained with DAPI (blue). Lower panels are zoomed in images of cells in white box in upper panels. **(B)** The number of PML nuclear bodies (NB) per cell was quantified in 54 cells per experiment for each infection condition. Each point represents the number of PML NB per cell. Lines represent the mean ± standard deviation. Graph shows a single representative experiment (n = 3). One-way ANOVA was performed using the mean for each biological replicate and comparing the sample to the the mock-infected control. The number of PML NB per cell was statistically lower than mock in SA11-4F-infected cells (p < 0.05), but not significantly different in SA11-5S- or OSU-infected cells. **(C)** The area of 100 PML NB was measured for each infection condition. Each point represents the area of a single PML NB. Line represents mean area ± standard deviation. Graph shows a single representative experiment (n = 3). One-way ANOVA was performed using the mean for each biological replicate and comparing the sample to the the mock-infected control. The area of PML NB per cell was statistically higher than mock in OSU-infected cells (p < 0.005), but not significantly different in SA11-4F- or SA11-5S-infected cells.

In SA11-4F-infected cells, the PML NB retained their round morphology (Fig. 5A), but there was a notable decrease in the number of PML NB per nucleus, to about half of the number found in mock-infected cells (Fig. 5B). The decrease in the number of PML NB in SA11-4F-infected cells was not accompanied by an increase in the area of PML NB (Fig. 5C). When PML protein was examined by immunoblot there was not a decrease in overall PML protein levels (data not shown), suggesting the possibility that SA11-like NSP1s may be causing a dispersal of PML NB, but are not inducing degradation of PML. PML NB in SA11-5S-infected cells retained their sphere-shaped morphology (Fig. 5A) and showed no change in number or area when compared to mock (Fig. 5B, 5C). In OSU-infected cells, PML NB took on an oblong appearance, suggesting a change in their morphology (Fig. 5A). This change was reflected by the near doubling of the average diameter of PML NB in OSU-infected cells to 0.56 μm^2^ (Fig. 5C). However, the number of PML NB in OSU-infected cells was similar to mock-infected cells (Fig. 5B), which suggests that the alteration in size and morphology was not due to fusion of existing PML NB.

### Changes to PML NB do not consistently group with SA11-4F-like or OSU-like NSP1s

NSP1 proteins can be generally grouped into SA11-like, based on their ability to induce IRF degradation, and OSU-like, based on their ability to induce β-TrCP degradation (24). To determine if changes to PML NB generally grouped with SA11-4F-like or OSU-like NSP1s, MA104 cells were infected with a panel of monoreassortant viruses that contain distinct NSP1 genes in the same genetic background of the SA11-L2 parental virus. The SNF and SRF viruses encode NSP1 proteins derived from K9 and RRV rotaviruses, respectively, and given their ability to induce IRF degradation are considered to have SA11-4F-like NSP1s (8, 20, 24). The SDF, SKF, and SOF viruses encode NSP1 proteins derived from DS-1, KU, and OSU rotaviruses, respectively, and are considered to have OSU-like NSP1s based on their ability to induce β-TrCP degradation (20, 24, 25). At 8 h p.i., cells were fixed and immunostained for the viral protein VP6 and PML, and nuclei were counterstained with DAPI.

The SA11-L2 virus is related to SA11-4F (the NSP1 proteins are identical), and as expected PML NB appeared morphologically similar in SA11-L2- and SA11-4F-infected cells (Fig. 6A). In addition, the number of PML NB in SA11-L2-infected cells was about half of that observed in mock-infected cells (Fig. 6B). Cells infected with the SA11-like viruses SNF and SRF also contained approximately 6 PML NB per cell, half of the number found in mock-infected cells. Like infection with OSU, the OSU-like viruses SOF and SKF resulted in somewhat larger and more oblong PML NB (Fig. 6A). While the OSU- and SOF-infected cells had approximately the same number of PML NB as mock-infected cells, there were nearly two-times the number of PML NB found in SKF-infected cells (Fig. 6B). Unexpectedly, the OSU-like virus SDF caused a reduction in the number of PML NB, which appeared morphologically similar to PML NB in cells infected with SA11-like viruses (Fig. 6B, 6A).

**FIG 6.**
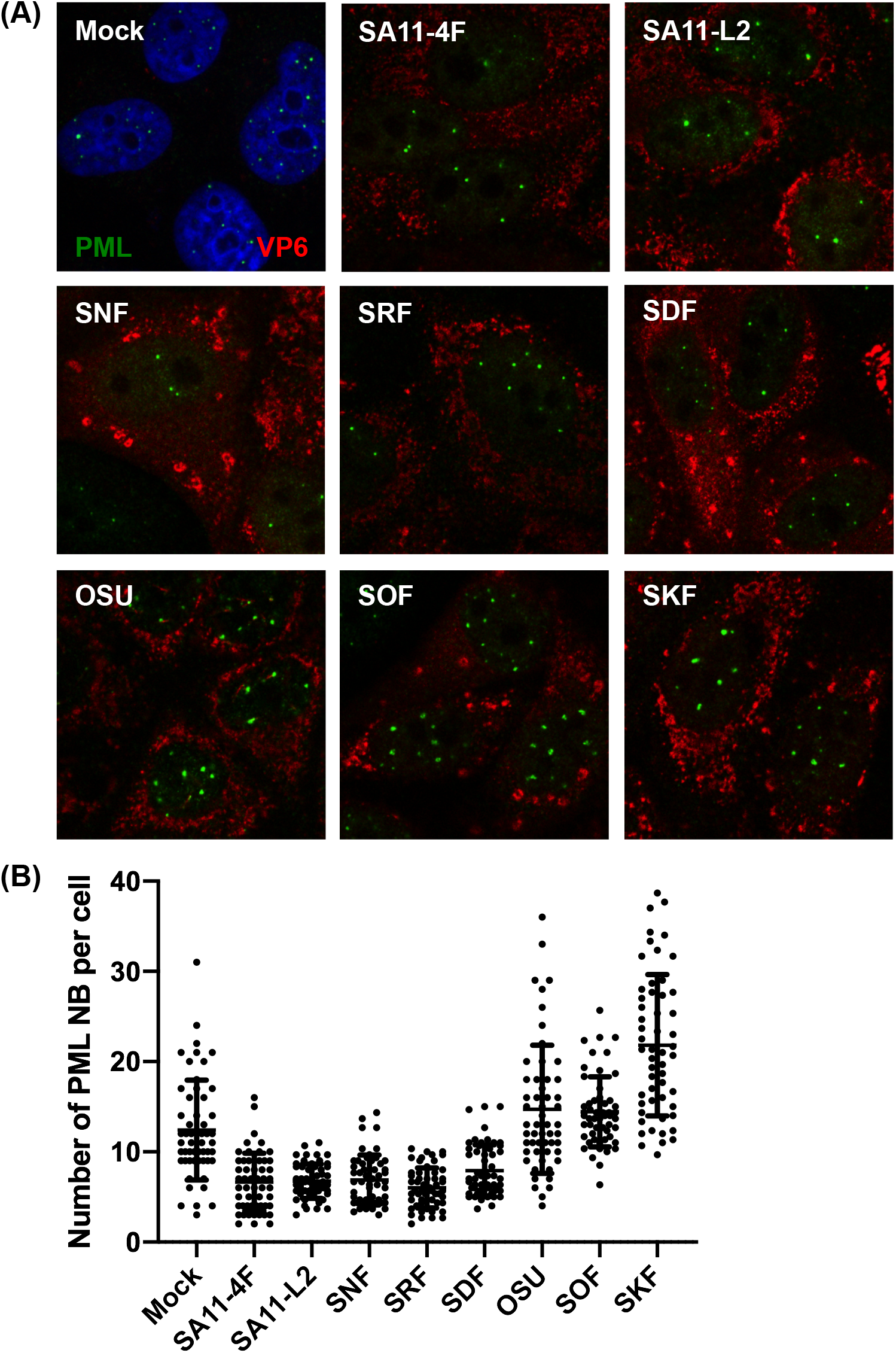
PML body morphology and quantity does not segregate into clear SA11-like or OSU-like groups. **(A)** MA104 cells were mock infected or infected with rotavirus strains as labeled (MOI = 5) for 8 h. Cells were fixed and stained with α-VP6 (red) and α-PML (green) antibodies followed by secondary staining with AlexaFluor594 goat α-guinea pig and AlexaFluor488 goat α-mouse. Nuclei were counterstained with DAPI (blue), but omitted for clarity in some images. **(B)** The number of PML nuclear bodies (NB) per cell was quantified in 54 cells per experiment for each infection condition. Each point represents the number of PML NB per cell. Lines represent the mean ± standard deviation. Graph shows a single representative experiment (n = 3). One-way ANOVA was performed using the mean for each biological replicate and comparing the sample to the mock-infected control. The number of PML NB per cell was statistically lower than mock in SA11-4F, SA11-L2, SNF, SRF, and SDF infected cells. The number of PML NB per cell was not statistically different than mock in OSU or SOF infected cells, but was statistically higher in SKF infected cells.

### OSU NSP1 nuclear foci no longer form in PML-deficient cells

To determine if PML is necessary for the formation of NSP1 foci in the nucleus of OSU-infected cells, HaCaT cells were transduced with lentivirus to knock down all isoforms of the PML protein by shRNA to create a stable cell line deficient in PML (shPML) (26). Previously published studies have shown no off-target effects using this same PML shRNA (26–30). HaCaT cells were also transduced with a lentivirus expressing a scrambled shRNA (shNEG) as a control (31). The loss of PML protein in shPML HaCaT cells was demonstrated by immunoblot, and quantification of PML protein levels consistently showed greater than 90% reduction (Fig. 7A). The loss of PML NB in shPML HaCaT cells was also evident by immunofluorescence microscopy (Fig. 7B). Next, the shPML and shNEG HaCaT cells were mock infected or infected with the OSU strain of rotavirus for 8 h followed by immunostaining for NSP1 and PML. In shNEG HaCaTs, the OSU NSP1 protein was diffusely distributed throughout the cytoplasm and found in distinct foci that colocalized with PML NB as had been observed in other cell types (Fig. 7C). In the shPML HaCaTs, the OSU NSP1 nuclear foci no longer formed in the absence of the PML protein, but some OSU NSP1 was still diffusely distributed in the nucleus. These results suggest that NSP1 nuclear localization is not dependent on the presence of PML, but that the formation of OSU NSP1 nuclear foci relies on the formation of PML NB.

**FIG 7.**
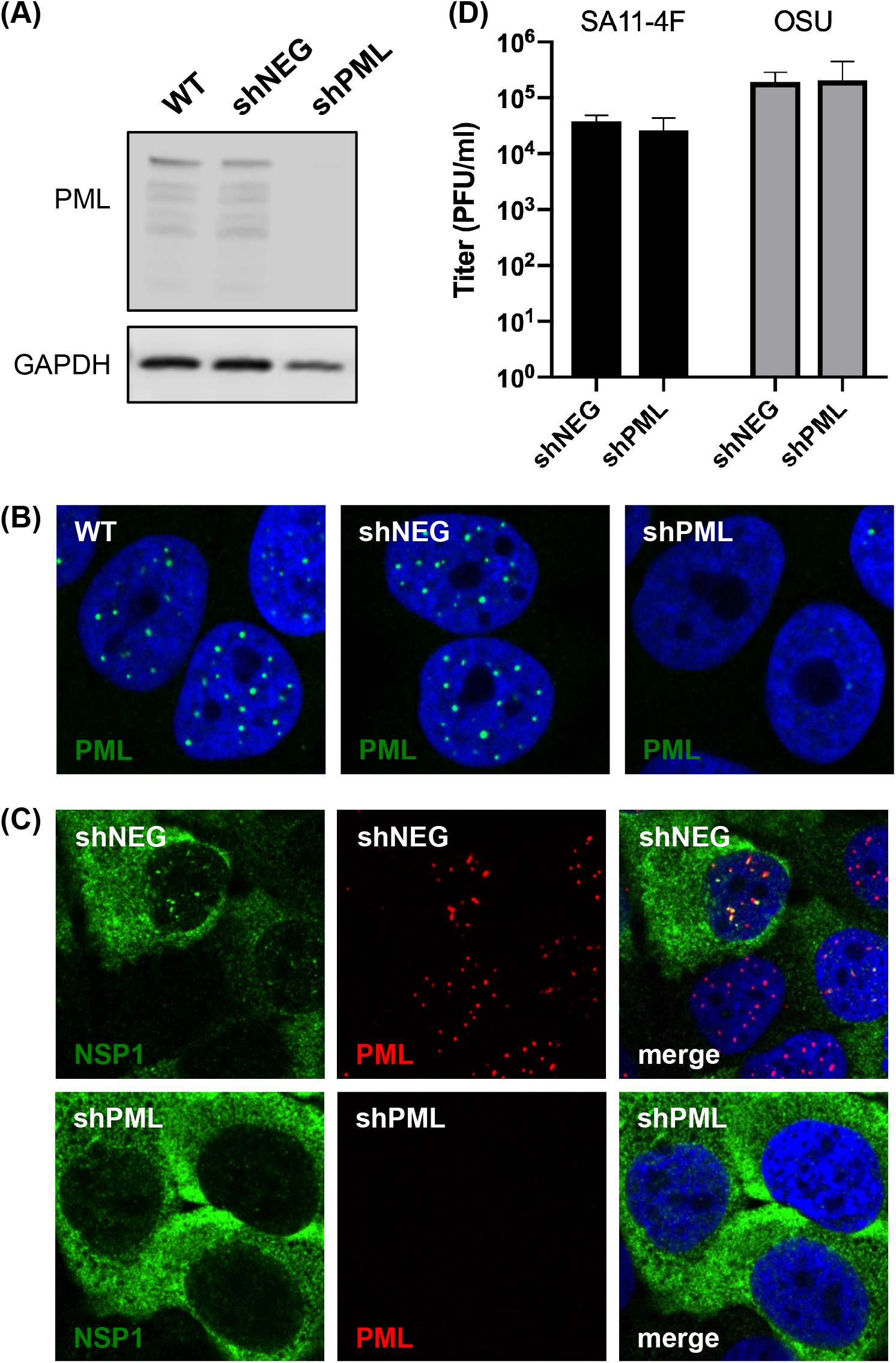
OSU NSP1 nuclear foci no longer form in PML-deficient cells. **(A)** HaCaT cells were transduced with shNEG (scrambled control) or shPML (PML knockdown) lentiviruses. Whole cell lysates were resolved by SDS-PAGE and analyzed by immunoblotting for PML or GAPDH (loading control). Blots were imaged using the Odyssey infrared imaging system (LiCor). **(B)** HaCaT cells were fixed and stained for PML (green) followed by secondary staining with AlexaFluor594 goat anti-mouse. Nuclei were counterstained with DAPI (blue). **(C)** shNEG and shPML HaCaT cells were infected with the OSU strain of rotavirus (MOI = 5) for 8 h. Cells were fixed and stained with α-OSU NSP1 (green) and α-PML (red) antibodies followed by secondary staining with AlexaFluor488 goat α-rabbit and AlexaFluor594 goat α-mouse. Nuclei were counterstained with DAPI (blue). (D) shNEG and shPML HaCaT cells were infected with the SA11-4F and OSU strains of rotavirus (MOI = 5) for 8 h. Cells were lysed by freeze-thaw and titered on MA104 cells.

To determine if the loss of PML affected viral titer, shNEG and shPML cells were mock infected or infected with SA11-4F or OSU rotavirus at a MOI of 5 for 8 h. Cells were lysed by multiple freeze-thaw cycles, and then viral titers were determined by plaque assay on MA104 cells. In shNEG HaCaT cells, the average titer of SA11-4F rotavirus was approximately 5.0 × 10^4^ PFU/ml, which was unchanged in shPML HaCaT cells (Fig. 7D). The average titer of OSU rotavirus was slightly higher at approximately 2.0 × 10^5^ PFU/ml in shNEG and shPML cells, but there was no measurable difference in OSU replication in the absence of PML protein. The importance of the association of OSU NSP1 with PML NB might not be adequately determined in highly permissive cell lines such as HaCaT and MA104, as it has previously been shown that rotaviruses expressing defective NSP1 proteins can replicate to similarly high titers as their wild-type parental counterparts in most cell lines (32–35).

## DISCUSSION

For DNA viruses that replicate in the nucleus of infected cells, there are a number of well-studied examples of viral genomes and proteins localizing to PML NB in infected cells (36). Some viruses with RNA genomes that replicate in the nucleus of infected cells have been shown to manipulate other types of nuclear bodies to enhance viral replication. There is now a growing appreciation for the manipulation of nuclear processes by cytoplasmically replicating viruses (37–40). Here we report that in addition to its known cytoplasmic localization, the rotavirus IFN antagonist protein NSP1 is also found in the nucleus of infected cells. Some rotavirus isolates, including OSU, WI61, and UK, formed punctate structures within the nucleus (Fig. 2). It was determined that the punctate NSP1 structures co-localized with PML NB (Fig. 4). Localization to the nucleus or to PML NB was not dependent on the C terminus of NSP1, which contains the substrate-binding domain that targets certain host proteins for proteasomal degradation (Fig. 2).

Previously published studies have concluded that NSP1 localizes to the cytoplasm of infected cells, based on immunofluorescence imaging (17–19). Re-examination of images from these studies indicates that NSP1 from SA11-5N, UK, UK variant brvA, and RRV did localize to the nucleus, sometimes as punctate spots, in addition to its cytoplasmic localization. Although the nuclear staining may have previously been dismissed as background, our use of subcellular fractionation confirms that NSP1 does indeed localize to the nucleus (Fig. 2). Given that the UK variant brvA expresses only the first 258 amino acids of the NSP1 protein due to the insertion of a premature stop codon, it is possible that the region responsible for nuclear localization or association with PML NBs is found in the N-terminal half of NSP1.

NSP1 has mainly been studied for its role in inhibition of the IFN-β response (reviewed in 41, 42) and host tropism (43, 44). SA11-4F-like rotaviruses have been shown to target IRFs for proteasomal degradation, whereas OSU-like rotaviruses have been shown to target β-TrCP for degradation (42). IRF or β-TrCP degradation prevents the activation of transcription factors that translocate to the nucleus, resulting in an inhibition of the type I IFN response. Given that the pathway of IFN induction begins in the cytoplasm and proceeds through a well-orchestrated series of signaling events, it is logical to predict that NSP1 is found only in the cytoplasm in order to induce IRF or β-TrCP degradation. However, IRFs and β-TrCP also localize to the nucleus; therefore, the role of NSP1 in the nucleus could potentially be to promote degradation of IRFs or β-TrCP in that cellular compartment. Transient expression of NSP1 proteins with small C-terminal truncations demonstrated that localization to the nucleus and PML NB occurred in the absence of the substrate binding domain, suggesting that the function of NSP1 in the nucleus is not related to degradation of IRFs or β-TrCP (Fig. 2).

PML NB have roles in a variety of cellular activities such as DNA damage and repair (45), apoptosis (46), and the IFN response (47–49). IFN-β induction leads to an increase in the number and size of PML NB (50, 51). Our data demonstrated that OSU infection caused an increase in the area but not the number of PML NB when compared to mock-infected cells (Fig. 5). Previous studies have shown that OSU NSP1 is able to inhibit IFN-β induction, although perhaps not as efficiently as the SA11-4F NSP1, which may account for the increase in PML NB area, but the reason for the size increase requires additional experimentation (8, 12). PML NB morphology is similarly altered in some DNA virus infections; for example, adenovirus E4 ORF3 causes PML NB to transform from sphere-shaped foci into track-like structures (52), and BK virus infection increases the size and decreases the number of PML NB to promote viral infection (53). Infection with other rotaviruses that express OSU-like NSP1 proteins (SDF, SOF, and SKF) did not consistently change the number of PML NB per cell; SDF infection caused a significant reduction, whereas SKF infection caused a significant increase in the number of PML NB (Fig. 6). While grouping NSP1 proteins into SA11-4F-like and OSU-like based on their target for degradation is convenient, it may not fully capture the spectrum of NSP1 activities in infected cells. Infection with other SA11-4F-like rotaviruses (SA11-L2, SNF, and SRF) resulted in a substantial reduction in the number of PML NB per cell when compared to mock, but the reason for such a reduction requires further exploration.

Many proteins other than PML reside in PML NB, and most often their localization is transient. Proteins that localize to PML NBs typically contain a SUMO-interacting motif (SIM) that is necessary for recruitment to sumolyated PML (54). SIM interactions with SUMO may favor the retention of proteins in PML NBs. At this time it is unknown if NSP1 associates with sumoylated PML or other NB-associated proteins, but studies are underway to identify a possible SIM motif in NSP1. It is thought that PML, global SUMO conjugation, and SUMO-dependent ubiquitination are tightly connected (55). NSP1 contains a conserved RING domain that bears similarity to other E3 ubiquitin ligases, and it is possible that the RING domain is essential for PML NB localization (23).

One of the challenges of studying NSP1 in highly permissive cell culture systems is the difficulty in ascertaining if the activities of NSP1 are essential to promote rotavirus replication, due to the fact that the NSP1 protein can be truncated or eliminated and the virus continues to replicate to high titers (17, 32–35). Although SA11-4F and OSU rotaviruses replicated to similar titers in cells lacking PML (Fig. 7D), the data is not sufficient to conclude that NSP1 localization to PML NB has no role in the viral life cycle. The development of a cell culture model that shows restricted replication in the absence of NSP1 is needed to better test the importance of NSP1 interactions with host proteins.

While this is the first report of a rotavirus nonstructural protein localizing to the nucleus of infected cells, NSP1 is not the first rotavirus protein shown to manipulate nuclear processes. The NSP3 protein expressed by rotaviruses, which binds to the eukaryotic translation initiation factor 4G and the 3’ consensus sequence of viral transcripts to alter translation, has been shown to disrupt nuclear-cytoplasmic transport of poly(A)-binding protein (PABP) (56). NSP3 appears to cause nuclear accumulation of poly(A)-containing mRNAs, thus preventing host mRNAs from reaching the cytoplasm to be translated (57). Other cytoplasmically replicating dsRNA viruses have also been shown to express proteins that localize to the nucleus. The reovirus μ2 protein, which interferes with the type I IFN response in a strain-specific manner (58, 59), has been shown to localize to nuclear speckles and alter host cell splicing (40). Orbiviruses including bluetongue virus (BTV) and African horse sickness virus express a nonstructural protein NS4 that has been shown to localize to the cytoplasm and nucleus of infected cells (60–62). BTV NS4, which also has been shown to antagonize the IFN response, is found in the nucleolus of infected cells but it does not appear to influence mRNA splicing or translation (63). Although each of these viruses and rotaviruses are members of the *Reoviridae* family, the structural and non-structural components vary widely. Though the mechanisms by which these viruses alter nuclear events to promote viral replication may vary, it is becoming clear that cytoplasmically replicating viruses utilize their limited coding space to modify the host and create a favorable replication environment.

## MATERIALS AND METHODS

### Cells and media

Human 293T cells were cultured in high glucose Dulbecco’s MEM (DMEM; Corning) supplemented with 5% fetal bovine serum (FBS) and 1% MEM non-essential amino acids (NEAA; HyClone). Human HT29 cells were cultured in high glucose DMEM supplemented with 10% FBS. Simian MA104 cells were cultured in Medium 199 (M199; Corning) supplemented with 5% FBS. Human HaCaT cells were cultured in low glucose Dulbecco’s MEM (DMEM; Corning) supplemented with 5% FBS. All cells were cultured at 37°C and 5% CO_2_.

### Viruses and infection

The rotavirus strains SA11-4F, SA11-5S (33), SA11-L2 (64), SDF, SKF, SNF (32), SOF, SRF (65), and OSU were propagated and quantified in MA104 cells as previously described (66). Viruses were activated by incubation with 10 μg/mL of trypsin for 30 min prior to infection of MA104 cells or 5 μg/mL of trypsin for 60 min prior to infection of HT29 or HaCaT cells. Cells were washed three times with serum-free medium and then inoculated with trypsin-activated rotavirus at a multiplicity of infection (MOI) of 5. At 1 hour post-infection (p.i.), cells were washed once with serum-free DMEM followed by addition of complete medium and incubation at 37°C for the specified time. Virus titers from infected HaCaT cell lysates were determined by plaque assay on MA104 cells as previously described (66).

### Antibodies

The following commercial antibodies and dilutions were used for immunofluorescence staining: rabbit polyclonal antibody to HaloTag (Promega; G9281; 1:500 dilution), mouse monoclonal antibody to lamin A/C (Cell Signaling; #4777; 1:150 dilution), peptide affinity purified rabbit polyclonal antisera to PML (Bethyl Laboratories; A301-167A; 1:500 dilution), mouse monoclonal antibody to SC-35 (Abcam; ab11826; 1:2,000 dilution), mouse monoclonal antibody to Gemin 2 (Abcam; ab6084; 1:250 dilution), and affinity purified rabbit polyclonal antibody to fibrillarin (CST Cat# cs-2639; 1:400 dilution). Affinity purified rabbit polyclonal antisera to NSP1 from SA11-4F and OSU were used at 1:500 dilution as previously described (8, 20). Polyclonal guinea pig antiserum to VP6 was used at 1:2,000 dilution as previously described (20). Secondary antibodies used for immunostaining were: AlexaFluor 488 goat anti-rabbit (Life Technologies; A-11034; 1:1,000 dilution), AlexaFluor 594 goat anti-mouse (Life Technologies; A-11032; 1:1,000 dilution), and AlexaFluor 594 goat anti-guinea pig (Life Technologies; A-11076; 1:1,000 dilution).

The following commercial antibodies and dilutions were used for immunoblotting: rabbit polyclonal antibody to HaloTag (Promega; G9281; 1:1,000 dilution), mouse monoclonal antibody to lamin A/C (Cell Signalling; #4777; 1:2,000 dilution), mouse monoclonal antibody to glyceraldehyde-3-phosphoate dehydrogenase (GAPDH) (Santa Cruz; sc-32233; 1:1,000 dilution), rabbit peptide affinity purified polyclonal antisera to PML (Bethyl Laboratories; A301-167A; 1:500 dilution), and affinity purified rabbit polyclonal antibody to calnexin (Cell Signaling; #2433; 1:2,000 dilution). Affinity purified rabbit polyclonal antisera to NSP1 from SA11-4F and OSU were used at 1:1,000 dilution as previously described (8, 20). Polyclonal antiserum to NSP2 from the simian SA11 strain of rotavirus was produced by Pacific Immunology Corporation (Ramona, CA). A peptide corresponding to amino acids 286 to 299 of SA11-5N NSP2 (C-KRLLFQKMKPEKNP) was conjugated to the carrier protein keyhole limpet hemocyanin. The peptide was used to immunize New Zealand White rabbits. NSP2-specific antiserum was collected and affinity purified using the immunizing peptide, which tested negative for cross-reactivity with other rotavirus proteins. Secondary antibodies used for immunoblotting were: IRDye 680RD goat anti-mouse (LiCor; 926-68070; 1:15,000 dilution), IRDye 800CW goat anti-rabbit (LiCor; 926-32211; 1:15,000 dilution), IRDye 680RD goat anti-rabbit (LiCor; 926-68073; 1:15,000 dilution), and IRDye 800CW goat anti-mouse (LiCor; 926-32210; 1:15,000 dilution).

### Plasmids and transfections

The following plasmids were used in transfection experiments: HaloTag vector (Promega; pHTN), pHTN-NSP1 (SA11-4F), pHTN-NSP1 (OSU), pHTN-NSP1 (WI61), pHTN-NSP1 (UK) or pHTN-NSP1 (DS-1) (20). To generate C-terminal truncations in the NSP1 proteins expressed from these plasmids, outward PCR amplification was used with each pHTN-NSP1 plasmid as a template and suitable primers. All plasmid sequences were verified by sequencing. For subcellular fractionation experiments, 3.0 × 10^6^ 293T cells were plated in 60-mm dishes, and transfected 24 h later with 2.5 μg of plasmid DNA with PolyJet (SignaGen, Rockville, MD). For immunofluorescence experiments, 5.0 × 10^5^ MA104 cells were plated in 6-well plates, and transfected the following day with 1.0 μg of plasmid DNA with PolyJet reagent. All transfection assays were analyzed at 48 h p.t.

### Subcellular fractionation

Cells were scraped into 1.5 mL cold phosphate buffered saline (PBS) and resuspended by vortexing. To generate whole cell lysates (WCL), 500 μL of cell suspension was centrifuged at 9.6 × *g* for 2 min to pellet cells. The supernatant was removed and the pellet was frozen at −80°C for at least 30 min. Cell pellets were lysed in 100 μL of radioimmunoprecipitation assay (RIPA) buffer (150 mM NaCl, 150 mM 1M Tris [pH 8.0], 0.5% sodium deoxycholate, 1.0% IGEPAL CA-630, 0.1% SDS, 1× protease inhibitor cocktail). Samples were incubated on ice for 20 min, then centrifuged at 18.8 × *g* for 3 min to pellet insoluble material. WCL was mixed with an equal part of 2× tricine sample buffer (100 mM 1M Tris-Cl [pH 6.8], 25% glycerol, 2% SDS, 0.02% bromophenol blue, 5% beta-mercaptoethanol) then boiled at 100°C for 5 min.

To generate the cytosolic fraction (CF) and nuclear fraction (NF), 1000 μL of cell suspension was centrifuged at 0.6 × *g* for 5 min to pellet cells. The supernatant was removed and pellet was resuspended in 150 μL of reticulocyte standard buffer (RSB) (10 mM Tris [pH 7.5], 16 mM NaCl, 3 mM MgCl_2_) containing 0.1 M sucrose, 1% IGEPAL CA-630, 0.5% sodium deoxycholate and 1 × protease inhibitor cocktail. Samples were incubated on ice for 15 min, then centrifuged at 0.9 × *g* for 5 min to separate the CF and NF. The CF was mixed with an equal part of 2× tricine sample buffer and boiled at 100°C for 5 min. The nuclear pellet was washed three times in 1 mL of PBS and stored at −80°C for at least 30 min. Nuclear pellets were lysed in 100 μL of RIPA buffer, incubated on ice for 20 min, and centrifuged at 9.6 × g for 2 min to pellet insoluble material. The NF was mixed with an equal part of 2× tricine sample buffer and boiled at 100°C for 5 min. Samples were stored at −80°C for long-term storage or used immediately for SDS-PAGE. All centrifugation steps were performed at +4°C.

### SDS-PAGE and immunoblotting

Proteins were resolved by SDS-PAGE in 10% tris-tricine gels and transferred to a nitrocellulose membrane (Li-Cor). Membranes were blocked in Odyssey TBS blocking buffer (Li-Cor) and then incubated with primary antibody in Odyssey blocking buffer containing 0.1% Tween 20. Membranes were washed three times with Tris buffered saline (50 mM Tris [pH 7.5], 150 mM NaCl) containing 0.1% Tween 20 (TBS-T). Secondary antibodies conjugated to IRDye680 or IRDye800 (Li-Cor) were added to TBS-T containing 1% nonfat dry milk. Membranes were washed three times with TBS-T and imaged using the Odyssey CLx infrared imaging system (Li-Cor) at a 5.5 intensity for each channel.

### Immunofluorescence staining and microscopy

Infected or transfected cells plated on glass coverslips were fixed by incubating with 11% formaldehyde in PBS at room temperature for 10 min. Fixed cells were permeabilized and blocked in PBS containing 0.05% Triton X-100 and 5% BSA at room temperature for 45 min. Permeabilized cells were incubated with primary antibody diluted in PBS containing 0.05% Triton X-100 and 1% BSA for 1 h at 37°C. Coverslips were washed four times in PBS at room temperature. Coverslips were then incubated with secondary antibody diluted in PBS containing 0.05% Triton X-100 and 1% BSA for 1 h at 37°C, followed by washing three times in PBS at room temperature. Coverslips were stained with 300 nM 4’,6-diamidino-2-phenylindole nucleic acid stain (DAPI; Invitrogen) in PBS for 5 min followed by washing three times in PBS at room temperature. Coverslips were rinsed briefly in sterile water and allowed to dry for 1 h prior to mounting to slides with ProLong Gold Antifade Reagent (ThermoFisher). All immunofluorescence images were captured using a Nikon SIM-E & A1R confocal microscope with an SR Apo TIRF 100x/1.49 NA oil objective lens. Images were processed uniformly with Adobe Photoshop software. The number and area of PML NB were quantified in maximum projection of z-stack images. The area of PML NB was calculated by manually drawing a polygonal structure around the edges of PML NB chosen at random using the Nikon NIS Elements software.

### Lentivirus transduction

Lentiviruses expressing anti-PML and anti control shRNAs were a kind gift from Dr. Roger Everett, MRC University of Glasgow Centre for Virus Research (26). The anti-PML shRNA coding strand sequence was 5’-AGATGCAGCTGTATCCAAG-3’, which lies in conserved exon 4. HaCaT cells were transduced with lentiviruses expressing shRNAs directed against a scrambled control (shNEG) or PML (shPML) by adding 1 mL of transduction media (low glucose DMEM, polybrene (5-8 ug/mL) and lentivirus) every 3 hours until the final volume was 3 mL. Cells were incubated for an additional 24 hours at 37°C prior to washing the transduction media and beginning selection with 10 ug/mL puromycin.

### Statistical analysis

For quantification of PML NB, the number per cell was counted in 54 cells per infection condition in a maximum intensity projection of confocal images. Counts were repeated in three independent experiments. One-way ANOVA analysis was performed using the mean number of PML NB per cell for each biological replicate and comparing the sample to the mean of the mock-infected control. For quantification of PML NB area, 100 PML NB were measured by manually drawing a polygonal shape around the edges randomly chosen PML bodies per infection condition in a maximum intensity projection of confocal images. Measurements were repeated in three independent experiments. One-way ANOVA analysis was performed using the mean area of PML NB per cell for each biological replicate and comparing the sample to the mean of the mock-infected control. Nikon NIS elements software was used for counting and area measurements. GraphPad Prism 8 was used for statistical analysis.

## ACKNOWLEDGEMENTS

Research reported in this publication was supported by an Institutional Development Award (IDeA) from the National Institute of General Medical Sciences of the National Institutes of Health under grant number P30GM110703 and by the Louisiana Board of Regents Research and Development Program under grant number LEQSF(2015-18)-RD-A-15. SKM was supported by an Ike Muslow Predoctoral Fellowship awarded by LSU Heath Sciences Center – Shreveport.

We are grateful to members of the Arnold lab for helpful discussions. We would like to thank Dr. Martin Sapp for generously providing cells and reagents, Dr. Malgorzata Bienkowska-Haba and Lucile Guion for their advice regarding reagents and confocal microscopy, Dr. Stephen DiGiuseppe for his assistance with immunostaining, and Dr. Martin Muggeridge for his assistance with subcellular fractionation.

## REFERENCES

1. Estes MK, Kapikian AZ. 2007. Rotaviruses, p. 1917–1974. In D. M. Knipe and P. M. Howley (ed.), Field’s Virology, 5th ed. Lippincott Williams & Wilkins, Philadelphia, PA.

2. Trask SD, McDonald SM, Patton JT. 2012. Structural insights into the coupling of virion assembly and rotavirus replication. Nat Rev Microbiol 10:165–177.

3. Silverman RH, Weiss SR. 2014. Viral phosphodiesterases that antagonize double-stranded RNA signaling to RNase L by degrading 2-5A. J Interferon Cytokine Res 34:455–463.

4. Zhang R, Jha BK, Ogden KM, Dong B, Zhao L, Elliott R, Patton JT, Silverman RH, Weiss SR. 2013. Homologous 2’,5’-phosphodiesterases from disparate RNA viruses antagonize antiviral innate immunity. Proc Nat Acad Sci USA 110:13114–13119.

5. Barro M, Patton JT. 2005. Rotavirus nonstructural protein 1 subverts innate immune response by inducing degradation of IFN regulatory factor 3. Proc Nat Acad Sci USA 102:4114–4119.

6. Feng N, Sen A, Nguyen H, Vo P, Hoshino Y, Deal EM, Greenberg HB. 2009. Variation in antagonism of the interferon response to rotavirus NSP1 results in differential infectivity in mouse embryonic fibroblasts. J Virol 83:6987–6994.

7. Holloway G, Truong TT, Coulson BS. 2009. Rotavirus antagonizes cellular antiviral responses by inhibiting the nuclear accumulation of STAT1, STAT2, and NF-kappaB. J Virol 83:4942–4951.

8. Arnold MM, Patton JT. 2011. Diversity of interferon antagonist activities mediated by NSP1 proteins of different rotavirus strains. J Virol 85:1970–1979.

9. Sen A, Rott L, Phan N, Mukherjee G, Greenberg HB. 2014. Rotavirus NSP1 protein inhibits interferon-mediated STAT1 activation. J Virol 88:41–53.

10. Sen A, Rothenberg ME, Mukherjee G, Feng N, Kalisky T, Nair N, Johnstone IM, Clarke MF, Greenberg HB. 2012. Innate immune response to homologous rotavirus infection in the small intestinal villous epithelium at single-cell resolution. Proc Nat Acad Sci USA 109:20667–20672.

11. Barro M, Patton JT. 2007. Rotavirus NSP1 inhibits expression of type I interferon by antagonizing the function of interferon regulatory factors IRF3, IRF5, and IRF7. J Virol 81:4473–4481.

12. Graff JW, Ettayebi K, Hardy ME 2009. Rotavirus NSP1 inhibits NFkappaB activation by inducing proteasome-dependent degradation of beta-TrCP: a novel mechanism of IFN antagonism. PLoS Pathog 5:e1000280.

13. Bagchi P, Nandi S, Nayak MK, Chawla-Sarkar M. 2013. Molecular mechanism behind rotavirus NSP1-mediated PI3 kinase activation: interaction between NSP1 and the p85 subunit of PI3 kinase. J Virol 87:2358–2362.

14. Nandi S, Chanda S, Bagchi P, Nayak MK, Bhowmick R, Chawla-Sarkar M. 2014. MAVS protein is attenuated by rotavirus nonstructural protein 1. PLoS One 9:e92126.

15. Nandi S, Chanda S, Bagchi P, Nayak MK, Bhowmick R, Chawla-Sarkar M, P. O. Staff. 2015. Correction: MAVS protein is attenuated by rotavirus nonstructural protein 1. PLoS One 10:e0131956.

16. Bhowmick R, Halder UC, Chattopadhyay S, Nayak MK, Chawla-Sarkar M. 2013. Rotavirus-encoded nonstructural protein 1 modulates cellular apoptotic machinery by targeting tumor suppressor protein p53. J Virol 87:6840–6850.

17. Silvestri LS, Taraporewala ZF, Patton JT. 2004. Rotavirus replication: plus-sense templates for double-stranded RNA synthesis are made in viroplasms. J Virol 78:7763–7774.

18. Hua J, Chen X, Patton JT. 1994. Deletion mapping of the rotavirus metalloprotein NS53 (NSP1): the conserved cysteine-rich region is essential for virus-specific RNA binding. J Virol 68:3990–4000.

19. Martinez-Alvarez L, Pina-Vazquez C, Zarco W, Padilla-Noriega L. 2013. The shift from low to high non-structural protein 1 expression in rotavirus-infected MA-104 cells. Memorias do Instituto Oswaldo Cruz 108:421–428.

20. Lutz LM, Pace CR, Arnold MM. 2016. Rotavirus NSP1 associates with components of the cullin RING ligase family of E3 ubiquitin ligases. J Virol 90:6036–6048.

21. Ding S, Mooney N, Li B, Kelly MR, Feng N, Loktev AV, Sen A, Patton JT, Jackson PK, Greenberg HB. 2016. Comparative proteomics reveals strain-specific beta-TrCP degradation via rotavirus NSP1 hijacking a host cullin-3-Rbx1 complex. PLoS Pathog 12:e1005929.

22. Ayala-Breton C, Arias M, Espinosa R, Romero P, Arias CF, Lopez S. 2009. Analysis of the kinetics of transcription and replication of the rotavirus genome by RNA interference. J Virol 83:8819–8831.

23. Arnold MM, Patton JT. 2009. Rotavirus antagonism of the innate immune response. Viruses 1:1035–1056.

24. Morelli M, Dennis AF, Patton JT. 2015. Putative E3 ubiquitin ligase of human rotavirus inhibits NF-kappaB activation by using molecular mimicry to target beta-TrCP. mBio 6.

25. Davis KA, Morelli M, Patton JT. 2017. Rotavirus NSP1 requires casein kinase II-mediated phosphorylation for hijacking of cullin-RING ligases. mBio 8.

26. Everett RD, Parada C, Gripon P, Sirma H, Orr A. 2008. Replication of ICP0-null mutant herpes simplex virus type 1 is restricted by both PML and Sp100. J Virol 82:2661–2672.

27. Everett RD, Murray J, Orr A, Preston CM. 2007. Herpes simplex virus type 1 genomes are associated with ND10 nuclear substructures in quiescently infected human fibroblasts. J Virol 81:10991–11004.

28. Sarkari F, Wang X, Nguyen T, Frappier L. 2011. The herpesvirus associated ubiquitin specific protease, USP7, is a negative regulator of PML proteins and PML nuclear bodies. PLoS One 6:e16598.

29. Smith MC, Box AC, Haug JS, Lane WS, Davido DJ. 2014. A phospho-SIM in the antiviral protein PML is required for its recruitment to HSV-1 genomes. Cells 3:1131–1158.

30. Sivachandran N, Wang X, Frappier L. 2012. Functions of the Epstein-Barr virus EBNA1 protein in viral reactivation and lytic infection. J Virol 86:6146–6158.

31. Bienkowska-Haba M, Luszczek W, Keiffer TR, Guion LGM, DiGiuseppe S, Scott RS, Sapp M. 2017. Incoming human papillomavirus 16 genome is lost in PML protein-deficient HaCaT keratinocytes. Cell Microbiol 19.

32. Okada J, Kobayashi N, Taniguchi K, Shiomi H. 1999. Functional analysis of the heterologous NSP1 genes in the genetic background of simian rotavirus SA11. Arch Virol 144:1439–1449.

33. Patton JT, Taraporewala Z, Chen D, Chizhikov V, Jones M, Elhelu A, Collins M, Kearney K, Wagner M, Hoshino Y, Gouvea V. 2001. Effect of intragenic rearrangement and changes in the 3’ consensus sequence on NSP1 expression and rotavirus replication. J Virol 75:2076–2086.

34. Tian Y, Tarlow O, Ballard A, Desselberger U, McCrae MA. 1993. Genomic concatemerization/deletion in rotaviruses: a new mechanism for generating rapid genetic change of potential epidemiological importance. J Virol 67:6625–6632.

35. Taniguchi K, Kojima K, Urasawa S. 1996. Nondefective rotavirus mutants with an NSP1 gene which has a deletion of 500 nucleotides, including a cysteine-rich zinc finger motif-encoding region (nucleotides 156 to 248), or which has a nonsense codon at nucleotides 153-155. J Virol 70:4125–4130.

36. Komatsu T, Nagata K, Wodrich H. 2016. The role of nuclear antiviral factors against invading DNA viruses: the immediate gate of incoming viral genomes. Viruses 8.

37. Goertz GP, McNally KL, Robertson SJ, Best SM, Pijlman GP, Fros JJ. 2018. The methyltransferase-like domain of chikungunya virus nsP2 inhibits the interferon response by promoting the nuclear export of STAT1. J Virol 92:e01008–18.

38. Liu YC, Kuo RL, Lin JY, Huang PN, Huang Y, Liu H, Arnold JJ, Chen SJ, Wang RY, Cameron CE, Shih SR. 2014. Cytoplasmic viral RNA-dependent RNA polymerase disrupts the intracellular splicing machinery by entering the nucleus and interfering with Prp8. PLoS Pathog 10:e1004199.

39. Kumar A, Buhler S, Selisko B, Davidson A, Mulder K, Canard B, Miller S, Bartenschlager R. 2013. Nuclear localization of dengue virus nonstructural protein 5 does not strictly correlate with efficient viral RNA replication and inhibition of type I interferon signaling. J Virol 87:4545–4557.

40. Rivera-Serrano EE, Fritch EJ, Scholl EH, Sherry B. 2017. A cytoplasmic RNA virus alters the function of the cell splicing protein SRSF2. J Virol 91:e02488–16.

41. Arnold MM, Sen A, Greenberg HB, Patton JT. 2013. The battle between rotavirus and its host for control of the interferon signaling pathway. PLoS Pathog 9:e1003064.

42. Morelli M, Ogden KM, Patton JT. 2015. Silencing the alarms: Innate immune antagonism by rotavirus NSP1 and VP3. Virology 479–480:75-84.

43. Broome RL, Vo PT, Ward RL, Clark HF, Greenberg HB. 1993. Murine rotavirus genes encoding outer capsid proteins VP4 and VP7 are not major determinants of host range restriction and virulence. J Virol 67:2448–2455.

44. Feng N, Yasukawa LL, Sen A, Greenberg HB. 2013. Permissive replication of homologous murine rotavirus in the mouse intestine is primarily regulated by VP4 and NSP1. J Virol 87:8307–8316.

45. Dellaire G, Bazett-Jones DP. 2004. PML nuclear bodies: dynamic sensors of DNA damage and cellular stress. BioEssays: news and reviews in molecular, cellular and developmental biology 26:963–977.

46. Krieghoff-Henning E, Hofmann TG. 2008. Role of nuclear bodies in apoptosis signalling. Biochimica et biophysica acta 1783:2185–2194.

47. El Asmi F, Maroui MA, Dutrieux J, Blondel D, Nisole S, Chelbi-Alix MK. 2014. Implication of PMLIV in both intrinsic and innate immunity. PLoS Pathog 10:e1003975.

48. Everett RD, Chelbi-Alix MK. 2007. PML and PML nuclear bodies: implications in antiviral defence. Biochimie 89:819–830.

49. Tavalai N, Stamminger T. 2008. New insights into the role of the subnuclear structure ND10 for viral infection. Biochim Biophys Acta 1783:2207–2221.

50. Chelbi-Alix MK, Pelicano L, Quignon F, Koken MH, Venturini L, Stadler M, Pavlovic J, Degos L, de The H. 1995. Induction of the PML protein by interferons in normal and APL cells. Leukemia 9:2027–2033.

51. Lavau C, Marchio A, Fagioli M, Jansen J, Falini B, Lebon P, Grosveld F, Pandolfi PP, Pelicci PG, Dejean A. 1995. The acute promyelocytic leukaemia-associated PML gene is induced by interferon. Oncogene 11:871–876.

52. Carvalho T, Seeler JS, Ohman K, Jordan P, Pettersson U, Akusjarvi G, Carmo-Fonseca M, Dejean A. 1995. Targeting of adenovirus E1A and E4-ORF3 proteins to nuclear matrix-associated PML bodies. J Cell Biol 131:45–56.

53. Jiang M, Entezami P, Gamez M, Stamminger T, Imperiale MJ. 2011. Functional reorganization of promyelocytic leukemia nuclear bodies during BK virus infection. mBio 2:e00281–00210.

54. Sahin U, Ferhi O, Jeanne M, Benhenda S, Berthier C, Jollivet F, Niwa-Kawakita M, Faklaris O, Setterblad N, de The H, Lallemand-Breitenbach V. 2014. Oxidative stress-induced assembly of PML nuclear bodies controls sumoylation of partner proteins. J Cell Biol 204:931–945.

55. Lallemand-Breitenbach V, de The H. 2018. PML nuclear bodies: from architecture to function. Curr Opin Cell Biol 52:154–161.

56. Arnold MM, Brownback CS, Taraporewala ZF, Patton JT. 2012. Rotavirus variant replicates efficiently although encoding an aberrant NSP3 that fails to induce nuclear localization of poly(A)-binding protein. J Gen Virol 93:1483–1494.

57. Rubio RM, Mora SI, Romero P, Arias CF, Lopez S. 2013. Rotavirus prevents the expression of host responses by blocking the nucleocytoplasmic transport of polyadenylated mRNAs. J Virol 87:6336–6345.

58. Irvin SC, Zurney J, Ooms LS, Chappell JD, Dermody TS, Sherry B. 2012. A single-amino-acid polymorphism in reovirus protein mu2 determines repression of interferon signaling and modulates myocarditis. J Virol 86:2302–2311.

59. Zurney J, Kobayashi T, Holm GH, Dermody TS, Sherry B. 2009. Reovirus mu2 protein inhibits interferon signaling through a novel mechanism involving nuclear accumulation of interferon regulatory factor 9. J Virol 83:2178–2187.

60. Zwart L, Potgieter CA, Clift SJ, van Staden V. 2015. Characterising non-structural protein NS4 of African horse sickness virus. PLoS One 10:e0124281.

61. Belhouchet M, Mohd Jaafar F, Firth AE, Grimes JM, Mertens PP, Attoui H. 2011. Detection of a fourth orbivirus non-structural protein. PLoS One 6:e25697.

62. Ratinier M, Caporale M, Golder M, Franzoni G, Allan K, Nunes SF, Armezzani A, Bayoumy A, Rixon F, Shaw A, Palmarini M. 2011. Identification and characterization of a novel non-structural protein of bluetongue virus. PLoS Pathog 7:e1002477.

63. Ratinier M, Shaw AE, Barry G, Gu Q, Di Gialleonardo L, Janowicz A, Varela M, Randall RE, Caporale M, Palmarini M. 2016. Bluetongue Virus NS4 Protein Is an Interferon Antagonist and a Determinant of Virus Virulence. J Virol 90:5427–5439.

64. Taniguchi K, Nishikawa K, Kobayashi N, Urasawa T, Wu H, Gorziglia M, Urasawa S. 1994. Differences in plaque size and VP4 sequence found in SA11 virus clones having simian authentic VP4. Virology 198:325–330.

65. Mahbub Alam M, Kobayashi N, Ishino M, Naik TN, Taniguchi K. 2006. Analysis of genetic factors related to preferential selection of the NSP1 gene segment observed in mixed infection and multiple passage of rotaviruses. Arch Virol 151:2149–2159.

66. Arnold M, Patton JT, McDonald SM. 2009. Culturing, storage, and quantification of rotaviruses. Curr Protoc Microbiol Chapter 15:Unit 15C 13.

